# Neuron-specific epigenetic repression of *Cdk5* impairs hippocampal-dependent memory in male and female mice

**DOI:** 10.64898/2026.04.08.716689

**Authors:** Kiara L. Rodríguez-Acevedo, Julia J. Winter, Megan I. Alvarez, Ajinkya Sase, Kyle Czarnecki, Elizabeth A. Heller

**Affiliations:** Department of Systems Pharmacology and Translational Therapeutics, University of Pennsylvania, Philadelphia, PA 19104, USA

## Abstract

Biological sex regulates fundamental neurobiology, as well as the etiology and prevalence of neuropsychiatric disorders. Cyclin-dependent kinase 5 (Cdk5) is a neuronally enriched kinase that regulates synaptic plasticity, neuronal homeostasis, and hippocampal-dependent memory. While Cdk5 protein activity is necessary and sufficient to promote memory in male rodents, its role in females and its gene regulation in either sex remain poorly understood. In males, Cdk5 protein inhibition impairs fear memory. We previously showed that fear conditioning activates *Cdk5* gene expression and increases permissive chromatin acetylation in male, but not female hippocampus. We hypothesize that *Cdk5* gene repression would impair fear memory in males. We developed an excitatory neuron-specific, CRISPR/dCas9-HDAC3 epigenetic editing tool to target histone acetylation at the endogenous Cdk5 promoter. This strategy reduced histone acetylation and decreased Cdk5 mRNA, protein, and kinase activity in both sexes. Interestingly, *Cdk5* repression in hippocampal neurons impaired fear and spatial memory in both male and female mice. Targeted deacetylation also evicted the transcription factor CREB1 from the Cdk5 promoter, revealing a link between histone acetylation and Cdk5 transcriptional activation. These findings demonstrate that *Cdk5* acetylation in neurons is necessary for hippocampal memory in both sexes, providing new insight into sex-specific epigenetic regulation of memory.

## Introduction

Sex differences in the brain emerge during development and shape neurophysiology, circuit function, and behavior across the lifespan ^1–3^. At the molecular level, sex differences in gene expression, epigenetic regulation, and synaptic plasticity have been documented across brain regions ^4–7^ yet the functional consequences of these differences for cognition and memory remain incompletely understood. When examining the neurobiology of learning specifically, sex differences in performance have been observed across numerous learning tasks in both human and literature^8^, suggesting that the neural mechanisms supporting cognition, memory acquisition and retrieval are not identical between sexes. These differences in fundamental neurobiology have important translational implications. Sex differences exist in the vulnerability and manifestation of neurological and psychiatric disorders involving cognitive disruption. Women are disproportionately affected with Alzheimer’s Disease (AD), while men show greater rates of schizophrenia^9^. Psychiatric disorders including Post-traumatic stress disorder (PTSD) and depression are twice more likely to be diagnosed in women^10–14^. Despite this, females have been historically excluded from neuropsychiatric preclinical models^15,16^, leaving the molecular and epigenetic mechanisms underlying these sex differences largely unknown.

Cyclin-dependent kinase 5 (Cdk5) is a non-canonical Cdk, enriched in neurons, that functions as a key regulator of synaptic plasticity, neuronal homeostasis, and hippocampal-dependent memory ^17,18^. Over three decades of research have established roles for Cdk5 protein activity in neuronal development, stress response, drug reward, and learning^19–25^, with these effects documented predominantly in males only. Altogether, these works point to an important role of Cdk5 in regulating learning and memory processes. Cdk5 phosphorylates Tau protein at multiple residues important for normal neurobiological function^26^, contributing to cytoskeletal dynamics and synaptic plasticity, but Cdk5-mediated hyperphosphorylation at pathological residues promotes Tau dissociation from microtubules, accumulation into neurofibrillary tangles, and downstream memory impairments^26,27^. This pathological Tau hyperphosphorylation has implicated Cdk5 in neurodegenerative diseases including AD^28–31^. Notably, female AD brains show markedly higher Tau phosphorylation at Cdk5-targeted residues compared to males^32^, suggesting sex-divergent Cdk5 regulation may contribute to differential vulnerability. Nevertheless, female subjects are largely excluded from studies on Cdk5 regulation in the brain, and sex as a biological variable is rarely formally analyzed.

Cdk5 protein activation is necessary for memory formation in males^33^ while its acute activation is sufficient to enhance hippocampal-dependent memory^34^, yet little is known on the role of *Cdk5* gene activation in either sex. *Cdk5* activation is regulated through dynamic histone acetylation ^35–37^. We previously used epigenetic editing to acetylate and activate *Cdk5*, finding this sufficient to impair fear memory in female but not male mice^37^. In the current study, we tested the necessity of *Cdk5* gene activation for fear conditioning in both sexes using CRISPR/dCas9-HDAC3 epigenetic editing^38^ to deacetylate and repress *Cdk5* gene expression specifically in excitatory neurons of the male and female mouse hippocampus. Unexpectedly, we found that *Cdk5* gene repression impaired hippocampal-dependent fear and spatial memory in both sexes. This finding helps address a critical gap in our understanding of how histone acetylation at a single gene locus shapes hippocampal memory in both sexes, with implications for sex-differences in the prevalence and severity of neuropsychiatric disorders.

## Results

### Epigenetic editing with CRISPR/dCas9-HDAC3 represses *Cdk5* expression *in vitro* and *in vivo*

Dynamic histone acetylation is observed at the *Cdk5* promoter in response to neuronal activation^35–37^ yet its direct causal relevance to *Cdk5* gene activation is not well understood. To address this, we developed a CRISPR/dCas9-HDAC3 epigenetic editing tool to remove endogenous histone H3K9/14 acetylation at a select genomic locus using a single-guide RNA (sgRNA) (Fig. 1a). We profiled histone H3K9/14 acetylation proximal to the *Cdk5* transcription start site (TSS) in naïve mouse hippocampus and designed five *Cdk5* sgRNAs that targeted an endogenous peak of *Cdk5* H3K9/14 acetylation 62 to 1107 bp upstream of the *Cdk5* TSS (Fig. 1b). We validated the epigenetic editing approach first in N2a cells, finding reduced Cdk5 mRNA by dCas9-HDAC3 + *Cdk5*-sgRNA-691 (henceforth referred to as *Cdk5*-sgRNA), compared to non-targeting (NT)-sgRNA control (Fig. 1c). *Cdk5*-sgRNA transfection with CRISPR/dCas9-AM, a control with no catalytic domain, produced no *Cdk5* repression (Fig. 1c), confirming that *Cdk5* gene repression was due to HDAC3 and not steric occlusion of the TSS by CRISPR/dCas9.

**Fig. 1.**
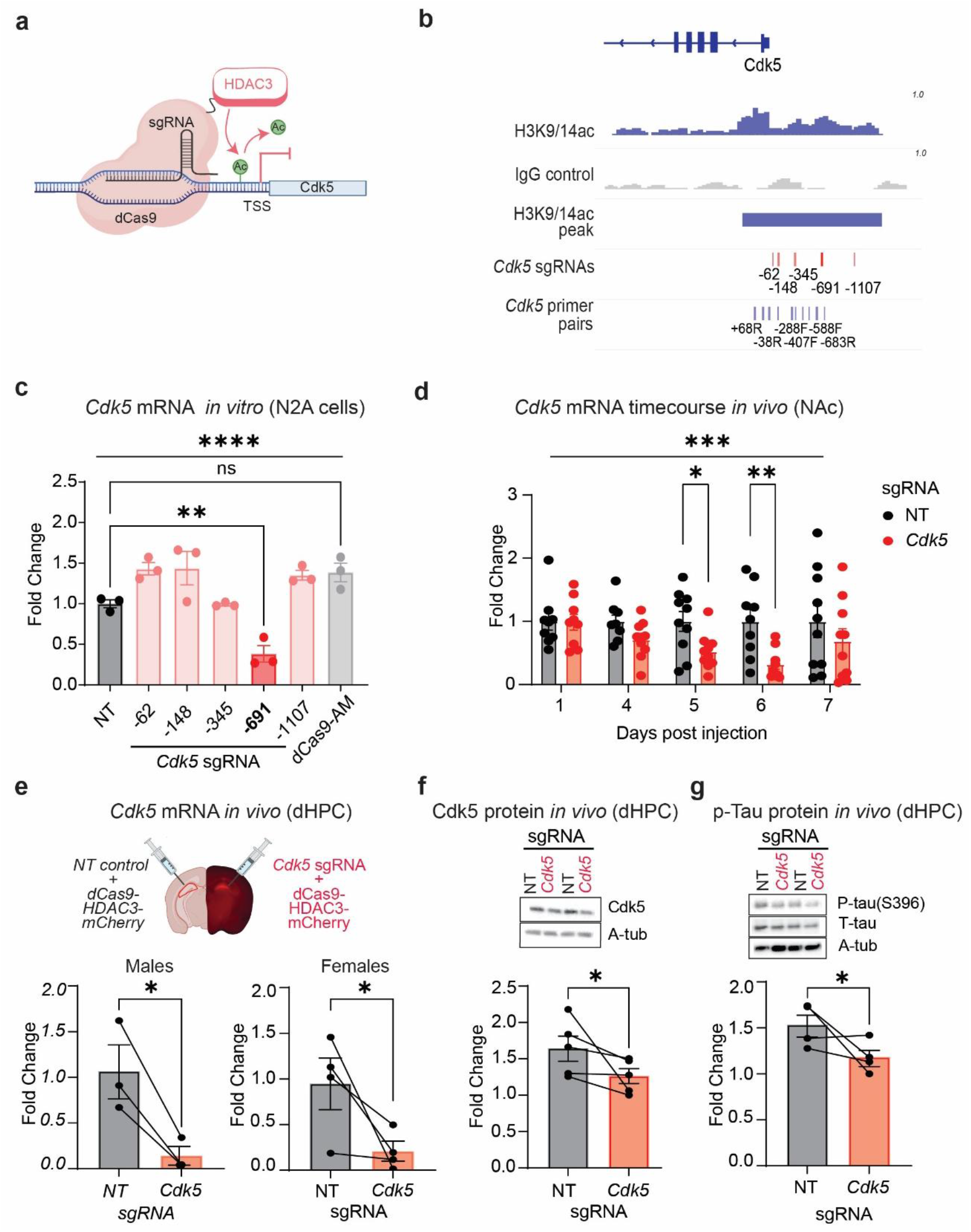
Targeted epigenetic repression of *Cdk5* with CRISPR/dCas9-HDAC3 reduces mRNA, protein and kinase activity in mouse brain. **(a)** Schematic of targeted histone deacetylation at the *Cdk5* promoter using CRISPR/dCas9-HDAC3. **(b)** CUT&Tag bigwig-normalized signal and called peak for H3K9/14ac in mouse hippocampus (top), and positions of designed sgRNAs targeting 62–1107 bp upstream of the *Cdk5* TSS (bottom). **(c)** *Cdk5* mRNA expression in N2a cells treated with dCas9-HDAC3 paired with each designed sgRNA, or dCas9-AM tag with *Cdk5*-sgRNA (*n* = 3). One-way ANOVA: main effect of sgRNA, F(6, 14) = 13.35, p < 0.0001; post-hoc NT vs. *Cdk5*-sgRNA, p = 0.0049; dCas9-AM tag vs. NT, p = 0.0907 (ns). **(d)** Intra-NAc injection of the neuronal hSyn-dCas9-HDAC3 with NT (left hemisphere) or *Cdk5*-sgRNA (right hemisphere) reduces *Cdk5* mRNA at days 5 and 6 post-injection (*n* = 9-10). Two-way ANOVA: main effect of sgRNA, F (1, 84) = 12.91, p = 0.0005; post-hoc NT vs. *Cdk5*-sgRNA at day 5, p = 0.0276; day 6, p = 0.0034. **(e)** Intra-dHPC injection of hSyn-dCas9-HDAC3 with NT (left hemisphere) or *Cdk5*-sgRNA (right hemisphere) reduces *Cdk5* mRNA at day 6 post-injection in males (left; paired t-test, p = 0.0395, *n* = 3) and females (right; paired t-test, p = 0.0425; *n* = 4). **(f)** hSyn-dCas9-HDAC3 with *Cdk5*-sgRNA reduces Cdk5 protein levels at day 6 PI in the dorsal hippocampus of a mixed-sex cohort (paired Wilcoxon test, p = 0.0312; *n* = 5). **(g)** hSyn-dCas9-HDAC3 with *Cdk5*-sgRNA reduces phosphorylation of Tau at Serine 396 in the dorsal hippocampus of a mixed-sex cohort (paired Wilcoxon test, p = 0.0292; *n* = 4). All data are presented as mean ± SEM.

We next expressed dCas-HDAC3 in the nucleus accumbens (NAc) under control of the human Synapsin (hSyn) promoter for neuronal-specific expression *in vivo*. We found that intra-NAc treatment with CRISPR/dCas9-HDAC3 + *Cdk5*-sgRNA reduced Cdk5 mRNA at 5 and 6 days post-injection relative to NT-sgRNA-treated contralateral controls in a mixed sex cohort (Fig.1d). We then asked whether this approach was effective in the hippocampus, a region critical for learning and memory^57,58^. Bilateral injection of hSyn-dCas9-HDAC3 + *Cdk5*-sgRNA into the dorsal hippocampus (dHPC) of male and female mice reduced Cdk5 mRNA at day 6 post injection (Fig. 1e). Cdk5 protein levels and kinase activity, as measured by depleted phosphorylation of Tau at Serine 396, a Cdk5 target residue^59^ were also reduced (Fig. 1f–g).

### Cre-dependent epigenetic editing with CRISPR/dCas9-HDAC3 depletes H3K9/14ac and represses *Cdk5* expression in excitatory pyramidal neurons

Because Cdk5 functions primarily as a neuronal kinase^19,60^ we probed its gene regulation specifically within excitatory hippocampal neurons using Cre-dependent (LS1L) expression of CRISPR/dCas9 in the NEX-Cre mouse transgenic line^39^ (Fig. 2a). LS1L-CRISPR/dCas9-HDAC3 expression in NEX-Cre+ hippocampus reduced *Cdk5* mRNA (Fig. 2b). We further ruled out off-target repression of a neighboring gene, *Slc4a2*, whose promoter is ∼990 bp from the *Cdk5*-sgRNA binding site (Fig. 2c). A significant Spearman correlation between H3K9/14ac enrichment and *Cdk5* mRNA fold change further supported that *Cdk5* acetylation levels are tightly associated with *Cdk5* transcription (Fig. 2d). Finally, we measured deacetylation across a region 68 bp downstream to 688 bp upstream of the *Cdk5* TSS. H3K9/14ac depletion was greatest at the *Cdk5* TSS and the *Cdk5*-sgRNA binding site (Fig. 2e). These data are consistent with prior work showing dCas9-HDAC3’s range of activity extends from its immediate sgRNA binding site to act most efficiently at the tail ends of endogenous histone acetylation enrichment^38^. Overall, we find that dCas9-HDAC3 paired with a rationally designed *Cdk5* guide RNA efficiently and specifically deacetylates *Cdk5* promoter histones and represses *Cdk5* expression in hippocampal excitatory pyramidal neurons.

**Fig. 2.**
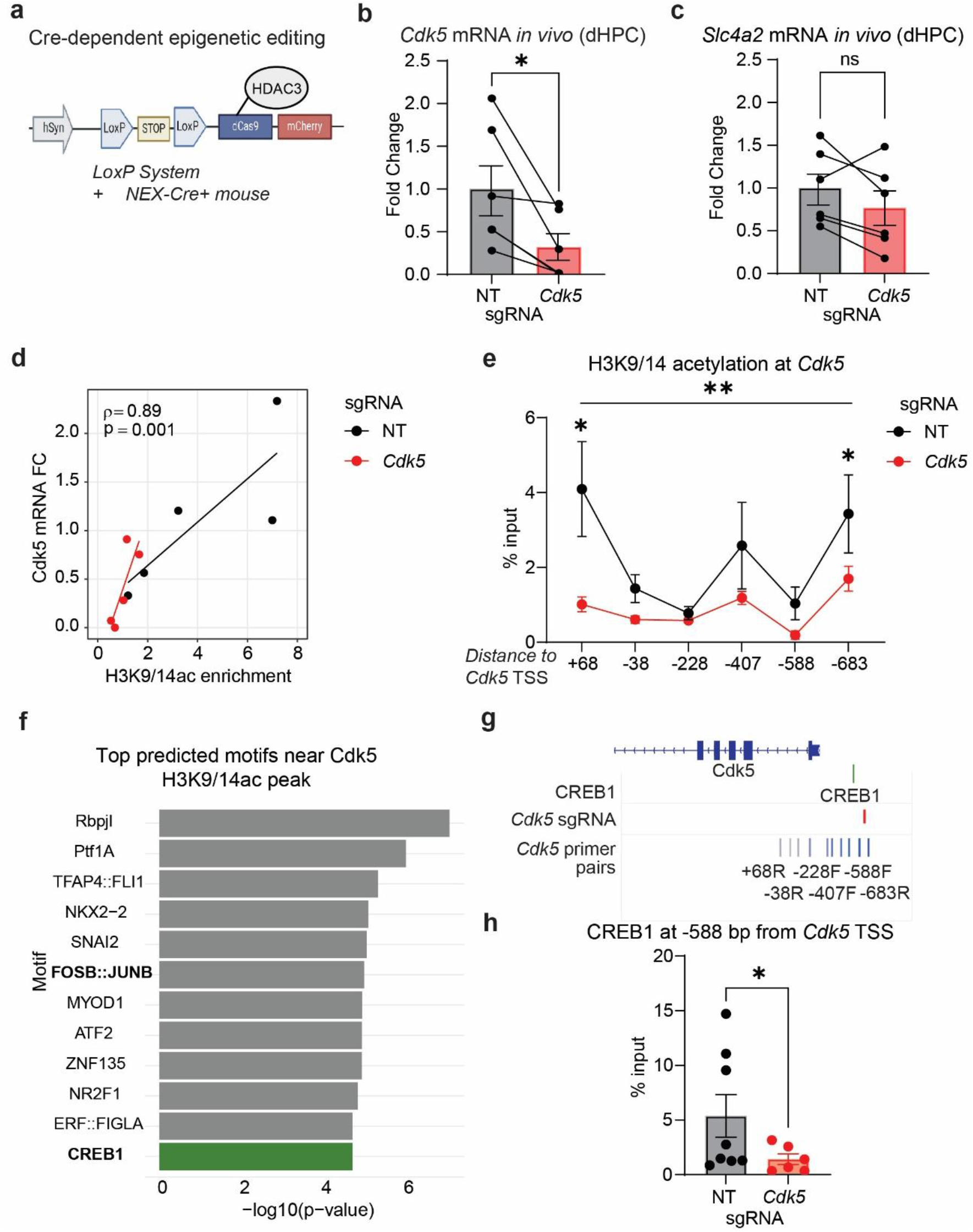
Excitatory neuron-specific histone deacetylation at *Cdk5* and neuronal loss of CREB1 occupancy following dCas9-HDAC3 editing. **(a)** Schematic of targeted histone deacetylation at the *Cdk5* promoter using LS1L-hSyn-dCas9-HDAC3, restricted to NEX-Cre+ neurons in mouse hippocampus. **(b)** Cell-type-specific histone deacetylation at the *Cdk5* promoter reduces *Cdk5* mRNA in NEX-Cre+ mice (paired t-test, p = 0.0283, *n* = 5). **(c)** Expression of *Slc4a2*, a gene with a promoter ∼990 bp from the *Cdk5*-sgRNA binding site, is not significantly affected (paired t-test, p = 0.1570, ns, *n* = 6). **(d)** H3K9/14ac enrichment positively correlates with *Cdk5* mRNA fold change across NT and *Cdk5*-sgRNA animals (Spearman *r* = 0.89, p = 0.001, *n* = 5). **(e)** H3K9/14ac enrichment measured by qChIP is reduced at primer pair 1 (midpoint ∼+68 bp from TSS) and primer pair 6 (midpoint ∼™683 bp from TSS). Two-way ANOVA: main effect of sgRNA, F(1, 47) = 12.79, p = 0.0008; main effect of primer pair, F(5, 47) = 3.940, p = 0.0046; post-hoc NT vs. *Cdk5*-sgRNA at +68 bp, p = 0.0113; at ™683 bp, p = 0.0410, *n* = 5). **(f)** MEME motif analysis of the endogenous H3K9/14ac peak in mouse hippocampus identifies FOSB:JUNB (TGTGAGGTCATC; score = 13.35, q = 3.73 × 10^−3^) and CREB1 (CTGTGAGGTCATC; score = 13.07, q = 7.12 × 10^−3^) as significantly predicted motifs. **(g)** The CREB1 motif (chr5:24,424,103-24,424,115) maps ∼588 bp upstream of the *Cdk5* TSS, within the region captured by the 588 bp primer pair. **(h)** CREB1 binding ∼588 bp upstream of the *Cdk5* TSS is reduced in *Cdk5*-sgRNA-treated animals as measured by qChIP (unpaired t-test with Welch’s correction, p = 0.0427; *n* = 8-9).

### Targeted histone deacetylation disrupts CREB1 binding at the *Cdk5* promoter

Histone deacetylation is associated with transcriptional repression through the loss of transcription factor recruitment^61^. Having linked histone deacetylation to *Cdk5* repression, we next investigated putative transcription factors at this locus. FosB binds *Cdk5* following cocaine exposure in the rat striatum^62^, and CREB1 directly binds the *Cdk5* promoter in rat brain^63^. Motif analysis on the *Cdk5* H3K9/14ac peak in mouse hippocampus predicted FosB:JunB and CREB1 (Fig. 2f), which share a binding motif approximately 588 bp upstream of the *Cdk5* TSS (Fig. 2g). We found that CREB1, but not FosB, was enriched at the predicted motif location (Fig. S2a–b), proximate to the site of maximal endogenous acetylation and depletion by dCas9-HDAC3 + *Cdk5-*sgRNA. We therefore asked whether H3K9/14ac depletion was sufficient to displace these factors (Fig. S2c). CRISPR/dCas9-HDAC3 *+ Cdk5*-sgRNA depleted CREB1 at the *Cdk5* promoter (Fig. 2h), compared to NT-sgRNA control. FosB binding did not change (Fig. S2d). We concluded that histone deacetylation at the *Cdk5* locus is sufficient to repress gene expression by evicting CREB1, but not FosB, from *the Cdk5* promoter in the mouse hippocampus.

### Epigenetic repression of *Cdk5* in excitatory neurons impairs hippocampal-dependent memory

We next asked whether epigenetic repression of *Cdk5* had functional consequences for memory. We injected LS1L-dCas9-HDAC3 + *Cdk5-* or NT-sgRNA into the hippocampus of NEX-Cre+ mice and performed contextual fear conditioning^37^ (Fig. 3a). Animals received three consecutive 0.4 mA foot shocks, and long-term memory retrieval was assessed 24 hours later. Animals treated with LS1L-dCas9-HDAC3+*Cdk5*-sgRNA exhibited reduced freezing during memory retrieval, compared to LS1L-dCas9-HDAC3+NT-sgRNA controls (Fig. 3b). Notably, *Cdk5*-sgRNA animals also displayed reduced freezing following the first and second shocks during acquisition, suggesting an initial delay in fear learning that resolved by the third shock. Control experiments confirmed that baseline fear behavior was comparable between NEX-Cre+ and -Cre™ animals across acquisition, 24-hour long-term recall, and 21-day remote recall (Fig. S3a), indicating that Cre expression alone does not affect freezing. Finally, there was no sex difference in the repression of long-term fear memory retrieval by *Cdk5* repression; a three-way ANOVA revealed a main effect of sgRNA and time, but no main effect of sex or significant interactions (Fig. S3b) while a two-way ANOVA revealed a significant *Cdk5*-sgRNA effect in both sexes (Fig. S3c–d).

**Fig. 3.**
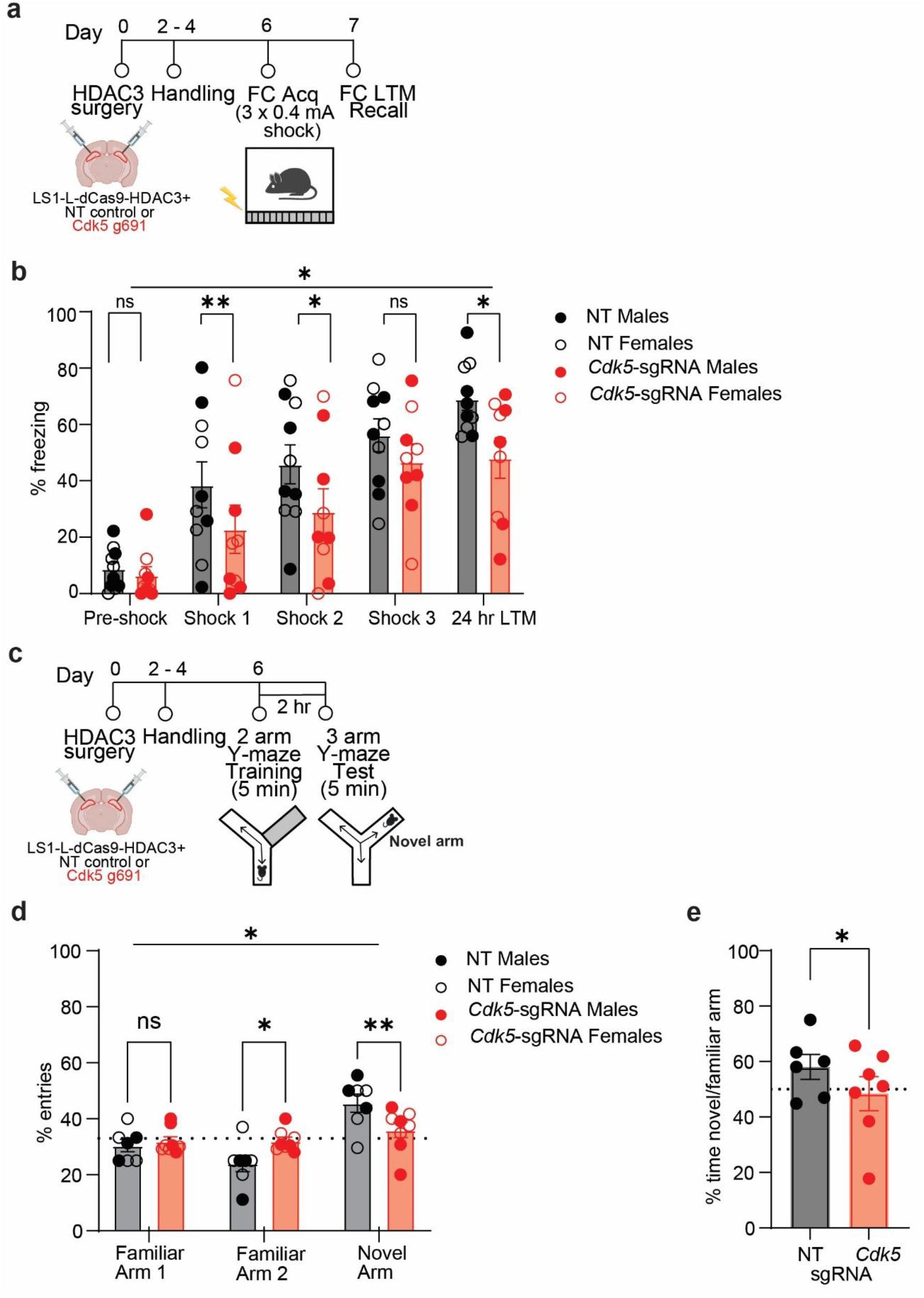
Epigenetic repression of *Cdk5* in hippocampal excitatory neurons impairs fear and spatial memory in males and females. **(a)** Experimental schematic of the multiple-shock fear conditioning protocol. At day 0, NEX-Cre+ animals received bilateral intra-dHPC injections of the Cre-dependent LSL-hSyn-dCas9-HDAC3 construct paired with NT or *Cdk5*-sgRNA. Following a 2-day recovery period and daily handling, animals underwent fear conditioning in a novel context and received three 0.4 mA foot shocks. Long-term memory was assessed 24 hours later in the same context. **(b)** *Cdk5*-sgRNA treated males and females exhibit reduced freezing compared to NT controls across the acquisition and retrieval phases (*n* = 8-10). Two-way ANOVA: main effect of sgRNA, F(1, 15) = 5.480, p = 0.0335; main effect of time, F(4, 59) = 49.34, p < 0.0001; sgRNA × time interaction, F(4, 59) = 2.716, p = 0.0382. Post-hoc comparisons: pre-shock, p = 0.7754 (n.s.); shock 1, p = 0.0028; shock 2, p = 0.0188; shock 3, p = 0.1632 (n.s.); 24-hour LTM recall, p = 0.0271. **(c)** Experimental schematic of the modified Y-maze reference memory task. **(d)** *Cdk5*-sgRNA-treated animals make fewer entries into the novel arm than NT controls (*n* = 7-8). Two-way ANOVA: main effect of arm, F(2, 39) = 15.41, p < 0.0001; sgRNA × arm interaction, F(2, 39) = 6.957, p = 0.0026. Post-hoc comparisons: NT vs. *Cdk5*-sgRNA novel arm entries, p = 0.0072; familiar arm entries, p = 0.0223. **(e)** *Cdk5*-sgRNA-treated animals spend less time in the novel arm than NT controls (unpaired t test, p= 0.0495). All data are presented as mean ± SEM.

To evaluate hippocampal-dependent spatial reference memory independently of aversive learning, we used a Y-maze reference memory task in which one arm was blocked during training and novel arm preference was measured at test (Fig. 3c). A three-way ANOVA revealed a significant main effect of arm and a sgRNA × arm interaction, with no significant effect of sex (Fig. S3e). *Cdk5*-sgRNA-treated animals of both sexes showed reduced entries to and time spent in the novel arm (Fig. 3d–e). Together, these findings demonstrate that histone deacetylation at the *Cdk5* promoter in excitatory hippocampal neurons is sufficient to impair both fear memory and spatial reference memory in male and female mice.

## Discussion

The role of Cdk5 kinase activity in neuronal function has been extensively characterized in both rodent and human brain^22,64,65^, with pharmacological inhibitors such as roscovitine traditionally used to interrogate the necessity of Cdk5 enzymatic activity in behavior and synaptic plasticity^66– 69^. Notably, prior work has demonstrated dose-dependent effects of Cdk5 activity on memory, wherein transient versus prolonged cofactor p25 expression produces enhancement versus impairment respectively^34^, underscoring that the precise level of Cdk5 activity is critical for normal memory function. Despite this sensitivity to Cdk5 activity levels, the transcriptional and epigenetic mechanisms regulating Cdk5 gene expression remain comparatively understudied. While the present study cannot fully disentangle the contributions of these distinct aspects of Cdk5 biology to the observed memory phenotypes, our epigenetic editing approach provides a complementary and precise tool for interrogating Cdk5 function at the level of gene regulation rather than protein activity.

The inclusion and analysis of females in our work further expand our mechanistic understanding of female fear memory, which is essential to understanding female biases in neuropsychiatric disease. Many prior studies demonstrate a role for *Cdk5* in hippocampal-dependent memory^17,18,70^, yet focused exclusively on males or did not specify sex in experimental approaches, therefore never formally analyzing sex as a biological variable. For example, conditional knockout of Cdk5 in pyramidal neurons leads to fear memory impairment in mice^17,70^. Here we discovered that targeted histone deacetylation and transcriptional repression of *Cdk5* in hippocampal pyramidal neurons impairs fear memory retrieval in both male and female mice. These findings are the first direct evidence that histone acetylation-dependent regulation of *Cdk5* in excitatory neurons contributes to hippocampal memory similarly in both sexes. Besides fear memory, targeted histone deacetylation at *Cdk5* also impaired hippocampal-dependent spatial memory in the Y-maze, extending the role of Cdk5 epigenetic regulation to non-aversive spatial memory. The convergence of impaired fear and spatial memory following a single epigenetic manipulation at the Cdk5 promoter suggests that histone acetylation at this locus is a broad regulator of hippocampal memory function rather than a fear-specific mechanism. This is consistent with the well-established role of Cdk5 in synaptic plasticity and LTP^24,71^, processes that underlie multiple forms of hippocampal-dependent memory.

The *in vivo* CRISPR-based epigenetic editing approach employed here expands beyond two prior tools: our previous zinc finger-based construct^36,37^, which was delivered in brain under a ubiquitous promoter, and the dCas9-HDAC3 system validated *in vitro* by Kwon et al.^38^. By driving expression under the neuron-specific hSyn promoter, the present tool restricts activity to neurons in the mouse brain and, when combined with Cre-dependent LSL constructs, enables cell-type-specific manipulation *in vivo*. Our histone deacetylation findings are consistent with Kwon et al., demonstrating de-enrichment at the tail ends of the endogenous acetylation peak. This consistency points to the importance of evaluating the endogenous chromatin landscape at the locus of interest, as the local acetylation profile may inform where effective histone modification is achievable. In our approach, we observe transcriptional repression of our target gene, *Cdk5*, 5-6 days post-injection. However, Cdk5 mRNA expression recovered by day 7 in the NAc, suggesting that epigenetic editing at this locus is temporally constrained. This recovery may reflect the re-establishment of endogenous acetylation by competing acetyltransferase activity, a decline in dCas9-HDAC3 expression or activity over time, or compensatory transcriptional mechanisms that restore Cdk5 expression despite sustained epigenetic editing. Whether similar recovery occurs in the dHPC remains to be determined, as our hippocampal time course was assessed at days 5-6 post-injection, likely capturing the window of maximal transcriptional repression.

The present findings extend the model proposed in our previous work^37^, in which females maintain a transcriptional brake on fear induced *Cdk5* activation to control for detrimental downstream effects. In that study, lifting this brake through *Cdk5* activation leads to fear memory impairment selectively in female mice, accompanied by female-specific tau hyperphosphorylation. The current findings add an important new dimension to this model: while females appear to actively suppress fear-induced Cdk5 transcription, baseline Cdk5 expression in excitatory hippocampal neurons is nonetheless required for normal memory function in both sexes. Together, these findings suggest that the female hippocampus is particularly sensitive to perturbations of Cdk5 expression in either direction. Future studies investigating the downstream targets of Cdk5 in excitatory neurons of both sexes, including tau and other synaptic substrates, will be critical for understanding the precise mechanisms through which *Cdk5* epigenetic dysregulation impairs hippocampal memory in a sex-dependent versus sex-independent manner.

Our finding that hSyn-dCas9-HDAC3-mediated *Cdk5* repression reduces tau phosphorylation at Serine 396 has important implications beyond fear memory. Cdk5 phosphorylation at tau Serine 396 is a hallmark of Alzheimer’s disease and pathologies^72–74^. Consistent with our findings, silencing Cdk5 via RNA interference reduces hippocampal tau neurofibrillary tangles in an Alzheimer’s Disease mouse model^28^. There are well-documented sex differences in tau pathology, in which females show greater vulnerability to tau hyperphosphorylation and faster progression of tau-related neurodegeneration than males in both human and preclinical studies^32,74–76^. Our current work suggests that epigenetic regulation of *Cdk5* is sufficient to modulate tau phosphorylation at a disease-relevant residue, positioning the Cdk5 promoter as a potential epigenetic target for reducing tau pathology in neurodegeneration.

Motif analysis of the H3K9/14ac peak near the Cdk5 TSS identified CREB1 and FosB as predicted transcriptional regulators at this locus, consistent with previous reports of CREB1 binding at the Cdk5 promoter in rat brain^63^ and FosB-dependent *Cdk5* activation in the rat striatum^23,62^. Notably, while FosB and CREB1 share a predicted binding motif in the *Cdk5* promoter, we found only CREB1 but not FosB enriched at this locus in mouse dorsal hippocampus, suggesting context-and region-specific occupancy of this regulatory element. Targeted histone deacetylation at the Cdk5 promoter was sufficient to evict CREB1 from this site, providing the first evidence linking H3K9/14ac enrichment to CREB1 occupancy at the Cdk5 locus. To our knowledge, this is the first demonstration that epigenetic remodeling at the *Cdk5* promoter directly regulates CREB1 transcription factor binding, adding a novel chromatin-level mechanism to the well-characterized role of CREB1 in hippocampal memory and fear-related gene expression^77–79^. Future studies examining whether CREB1 overexpression is sufficient to restore H3K9/14ac at the Cdk5 promoter, or whether CREB1 recruitment drives histone acetyltransferase activity at this locus, will be important for establishing the directionality of this relationship and the precise molecular hierarchy linking chromatin state to Cdk5 transcriptional regulation.

Together, these findings establish histone acetylation at the *Cdk5* promoter as a key epigenetic mechanism regulating excitatory neuron-dependent hippocampal memory in both sexes, with broad implications for understanding the molecular basis of fear-related psychiatric disorders and the potential development of locus-specific epigenetic interventions for their treatment.

## Materials and Methods

### Animals

10–16-week-old C57BL/6J male and female mice were purchased from Jackson Laboratory. 10–16-week-old Nex-Cre/+^39^; BirA/+ male and female mice were bred in-house on a C57BL/6 background. Male Nex-Cre/+ mice were paired with female BirA/+ mice to generate experimental (Nex-Cre/+; BirA/+) and control (Nex-Cre/-; BirA/+) offspring. All animals were housed in a temperature- and humidity-controlled vivarium under standard conditions. Pups remained in their birth cage until postnatal day 21 (P21), at which point they were weaned and housed in same-sex groups of 2–5 mice per cage. Mice remained in group housing through adulthood. Food and water were provided ad libitum throughout the experiment. Animals were maintained on a 12-hour (hr) light/dark cycle (lights on from 07:00 to 19:00). All behavioral testing was conducted during the light phase, between 07:00 and 18:00. All procedures were approved by and conducted in accordance with the guidelines of the Institutional Animal Care and Use Committee (IACUC) at the University of Pennsylvania.

### CRISPR and sgRNA design and synthesis

The dCas9-HDAC3 plasmid (Addgene, #98591)^38^ was subcloned into either an hSyn-IRES-GW-mCherry destination vector for pan-neuronal expression or an LS1-L-mCherry-GW destination vector for Cre-dependent, cell-type-specific expression (both gifts from Peter Hamilton) using Gateway cloning (LR Clonase II, Thermo Fisher Scientific). sgRNAs targeting the *Cdk5* promoter were designed within a window spanning ™62 to ™1107 bp upstream of the *Cdk5* transcription start site (TSS) using CRISPR-ERA^40^. sgRNA positions are reported as distance (bp) upstream *Cdk5* TSS (chr5:24,423,530), accounting for minus-strand orientation. Potential off-target effects were assessed computationally using Cas-OFFinder^41^. sgRNAs were synthesized as gBlocks (Integrated DNA Technologies), cloned into pENTR/dTOPO via BP recombination (Thermo Fisher Scientific, #K2420020), and subsequently transferred into the p1005 destination vector via LR recombination. All sgRNA constructs were verified by Sanger sequencing prior to use. For control experiments, a non-targeting sgRNA was designed with a scrambled sequence confirmed to have no complementarity with the mouse genome. To validate chromatin remodeling activity independent of dCas9 steric occlusion, a recombinant dCas9-AM tag fusion protein (Active Motif #81068, 100 ng) was used in combination with the sgRNAs *in vitro*.

### Single sample RNA and chromatin extraction

S3EQ cytoplasmic RNA and nuclear chromatin isolation from the same sample was performed as described previously^42^. Cytoplasmic RNA was purified using RNeasy Micro Kit (Qiagen) while nuclei were immediately stored at -80°C for downstream ChIP or CUT&TAG experiments.

### CUT&TAG-seq

CUT&Tag was performed according to published protocols^43^. Whole hippocampal hemispheres were hand dissected and flash frozen. Nuclei pellets were resuspended in Wash Buffer (20 mM HEPES-NaOH pH 7.5, 150 mM NaCl, 0.5 mM spermidine, 5 mM Sodium Butyrate and protease inhibitors). CUTANA Concanavalin A Conjugated Paramagnetic Beads (Epicypher) were activated in binding buffer before they were added to the nuclei suspension. Nuclei and beads were rotated for 10 min at 4°C and then anti-H3K9/14ac (Millipore, #17-615, 1:50 dilution) or IgG control (Novus Bio, #NBP2-24891, 1:50 dilution) antibodies were added to the mix at 4°C overnight with rotation. Bead-bound nuclei were washed 2x in digitonin buffer and pA-Tn5 adapter complex (Epicypher) was added for 1 hr. Tagmentation was stopped with SDS, EDTA and Proteinase K. DNA extraction was performed by phase separation using phenol:chloroform:isoamyl alcohol (25:24:1).

Libraries were prepared using the NEBNext® Ultra™ II DNA Library Prep Kit for Illumina (New England Biolabs). Adapters were ligated to DNA fragments, which were then amplified by PCR using a Universal i5 primer, uniquely barcoded i7 primers, and NEBNext High-Fidelity 2x PCR Master Mix. The PCR cycle consisted of an initial denaturation at 98°C for 30 s, followed by 13 cycles of 98°C for 10 s and 65°C for 10 s, and a final extension at 65°C for 5 min. PCR products were purified using 1.1x volume of Ampure Beads (Beckman Coulter) and washed with 80% ethanol. The cleaned DNA was eluted in 0.1X TE buffer. Library size distribution was confirmed using the Agilent TapeStation System (Agilent Technologies). Libraries were sequenced in paired-end mode on the Illumina NovaSeq X Plus platform, generating approximately 20 million reads per sample. Raw FASTQ files were processed and quality-checked using FastQC (v0.11.9)^44^. Adapter sequences were trimmed using BBduk (v38.84)^45^. Trimmed reads were aligned to the mm10 reference genome using Bowtie2 (v2.4.1)^46^. Uniquely mapped reads with a quality score >20 were retained using SAMtools (v1.13)^47^. Read alignments were normalized to total mapped reads using deepTools (v3.5.4.post1)^48^. Peaks were called using sicer2^49^ using IgG as a control.

### Motif Analysis

Motif enrichment analysis was performed using Simple Enrichment Analysis (SEA) from the MEME Suite^50^. The JASPAR CORE^51^ vertebrate motif database was used as the primary reference set of transcription factor motifs. Results were compared with additional curated motif collections, including the Mouse UniPROBE^52^ and Jolma 2013^53^ transcription factor motif databases. H3K9/14ac peaks identified by sicer2 were used as the input sequences. Control sequences were generated by shuffling the input peak sequences while preserving dinucleotide frequencies. SEA was run with a significance threshold of 0.05, and statistical significance was evaluated using false discovery rate (FDR)–adjusted *q*-values to account for multiple hypothesis testing. To identify transcription factor motifs specifically present at the Cdk5-associated H3K9/14ac peak, motif scanning was performed using Find Individual Motif Occurrences (FIMO) from the MEME Suite. The genomic sequence corresponding to the full Cdk5 peak was extracted and scanned against the JASPAR motif using a motif match threshold of *p* < 1 × 10^−4^. Motif occurrences were ranked by statistical significance and visualized using RStudio and the tidyverse package suite^54^.

### N2a Transfections

Murine Neuro-2a (N2a) cells (ATCC, #CCL-131) were maintained in EMEM (ATCC, #30-2003) supplemented with 10% fetal bovine serum (FBS; ATCC #30-2020). Cells were cultured at 37°C and 5% CO2 and passaged at 70-90% confluency using Trypsin/EDTA (ATCC #30-2101). Cryopreservation was performed in EMEM containing 10% FBS and 5% DMSO, with vials stored in liquid nitrogen. Cells were 70-80% confluent at the time of transfection with Effectene reagent (Quiagen). dCas9-HDAC3 (375 ng) and sgRNA (125 ng) were added to each well. As a control, dCas9-AM tag (Active Motif, #81068, 100 ng) was added instead. After 72 h, cells were collected and mRNA extracted as detailed above.

### Intracranial stereotaxic surgery

Stereotaxic intracranial infusions of plasmid DNA were performed as described previously^36,37,55^. Mice were anesthetized with isoflurane delivered via inhalation and secured in a stereotaxic apparatus (Kopf). Meloxicam (1:10 dilution) was administered subcutaneously at a dose of 2 µL per 20 g of body weight as a post-surgical analgesic. To target the NAc, we used +1.6 anterior/posterior, +/-1.5 medial/lateral, -4.4 dorsal/ventral coordinates relative to bregma at an angle of 10° from the midline. To target the dHPC we used -2.2 anterior/posterior, +/-2.0 medial/lateral, -1.8 dorsal/ventral relative to bregma at an angle of 7° from the midline. For plasmid transfections, jetPEI (Polyplus Sartorius, #89129-960) was used according to the manufacturer’s instructions. We delivered plasmid DNA at a concentration of 1 µg/µl in 1.5 µl jetPEI/plasmid solution per side and animal at a flow rate of 0.2 µl per minute, followed by 5 min rest before removing the syringes (Hamilton). After surgery, animals were allowed to recover for 3-5 days and daily monitored.

### Tissue collection

Animals were euthanized by rapid cervical dislocation. Whole brains were immediately extracted and submerged in ice-cold 1X phosphate-buffered saline (PBS; Sigma-Aldrich) supplemented with a protease inhibitor cocktail (cOmplete Protease Inhibitor Cocktail, Roche). Brains were then transferred to an adult mouse brain matrix (Zivic Instruments) and sectioned into 1 mm coronal slices. Bilateral 1.2–2 mm tissue punches (WellTech) were taken from the dorsal hippocampus and immediately frozen on dry ice. For animals receiving intra-NAc or intra-dHPC infusions of epigenetic editing CRISPR constructs, targeting accuracy and visual plasmid expression were confirmed by fluorescence microscopy using mCherry and GFP reporters (Leica). For ChIP experiments, two 1.2 mm punches per animal were pooled. All tissue samples were stored at ™80°C until downstream processing.

### Chromatin Immunoprecipitation (ChIP) and RT-PCR

Nuclei were fixed with 1% formaldehyde for 8–12 minutes and cross-linking was quenched with glycine. Chromatin was sheared using a Bioruptor, with 10% reserved as input control. Antibodies against H3K9/14ac (Millipore #17-615), CREB1 (Millipore #06-863), or FosB (Cell Signaling #2251) were bound to Dynabeads M280 Sheep anti-Rabbit (Thermo Fisher) for 6 hours, followed by overnight incubation with 700–900 µL sheared chromatin at 4°C. Bead-bound chromatin was washed, eluted, and reverse cross-linked with NaCl for 4 hours at room temperature. DNA was purified using the QIAamp Micro DNA Kit (Qiagen) and quantified by Qubit. ChIP enrichment was calculated by normalizing IP values to the adjusted percent input. For RNA, RT-PCR to generate cDNA was performed using iScript cDNA Synthesis Kit (Bio-Rad). The primer sequences are detailed in Supplementary Tables 1–2. Primers were named by distance (bp) between the amplicon midpoint, and the transcription start site (TSS), with sign indicating position relative to the TSS. For analysis of qRT-PCR, we used the DDCt method, normalizing to housekeeping genes *Gapdh* and *Rpl13a*.

### Protein Extraction and Western Blotting

One to two 1.2 mm tissue punches were homogenized in RIPA buffer (50mM Tris HCL, 150mM NaCl, 0.25% SDS, 0.25% Sodium Deoxycholate, 1mM EDTA) and supplemented with Halt Protease and Phosphatase Inhibitor Cocktail (Thermo Scientific) and incubated on ice for 10 minutes. Lysed cell homogenate was sonicated on ice at maximum intensity for 30 seconds and centrifuged at 14,000g at 4°C for 10 min. Supernatant protein was aliquoted and then quantified using Pierce BCA assay (Thermo Scientific) as per the manufacturer’s guidelines. For electrophoresis, 2-10 μg of each protein sample was reduced using 1x Lithium Dodecyl Sulphate buffer and 1x Sample Reducing Agent. The samples were then loaded onto a 4-20% SDS-PAGE gel (Bio-Rad) and electrophoresed at 90V for 1.5 hours or until the tracking dye reached the end of the gel. A molecular weight marker (Amersham Rainbow Marker-High range, GE Life Sciences) was included for reference. The separated proteins were subsequently transferred for 1.5 hours onto a Polyvinylidene difluoride membrane (Bio-Rad). The membrane was blocked with 5% Bovine Serum Albumin (Invitrogen Bioreagents) in 1x TBST (Tris-buffered saline with 0.1% Tween 20) for 1 hour at room temperature. Following the blocking step, the membranes were incubated overnight at 4°C with anti-Cdk5 antibody (Cell Signaling 2506S) at a 1:1000 dilution. The membranes were washed with 1x TBST and incubated with a secondary antibody, anti-rabbit IgG-HRP conjugated (Cell Signaling 7074) at a 1:3000 dilution for 1 hour at room temperature. The membranes were washed again with 1x TBST and imaged using an ECL substrate (Clarity Bio-Rad) and an Amersham imager 600 (GE Life Sciences). The immunoreactive bands were quantified in arbitrary units of optical density using the Image J software program (Fiji (imagej.net). After imaging, membranes were stripped (Restore Stripping Buffer 21059, Thermo Scientific) as per manufacturer’s guidelines and re-blotted with a-tubulin (Cell Signaling #2144S, 1:1000 dilution). Density was calculated by taking the ratio of optical density for Cdk5 and optical density for a-tubulin for each lane. Protein fold change was calculated by normalizing to the lowest optical density value.

### Contextual Fear Conditioning

Fear conditioning was performed in plexiglass chambers with wired floors (Med Associates, St. Albans, VT, USA) as described previously^37^. Each mouse was placed in the conditioning chamber and allowed to habituate for 148 s prior to shock delivery. Mice then received three 2 s foot shocks (0.4 mA), each separated by a 60 s inter-shock interval. Following a final 60 s period, mice were returned to their home cage. Long-term memory (LTM) was assessed 24 hours later by returning each mouse to the conditioning chamber for 5 min in the absence of shock. Freezing behavior during acquisition and retrieval was recorded and scored using VideoFreeze software (Med Associates). Freezing was defined as the absence of movement for ≥2 s, excluding respiratory motion, and is expressed as a percentage of total time in the chamber. The chamber was cleaned with 70% ethanol between subjects and cleaned twice between sex cohorts.

### Y-maze

A modified Y-maze spatial reference memory task was performed as described previously^56^. The Y-maze apparatus (Harvard Bioscience) consisted of three identical arms arranged in a “Y” configuration. During the acquisition (training) phase, one arm was blocked with a guillotine door. Each mouse was placed in one of the two accessible arms with its head oriented away from the center of the maze and allowed to freely explore for 5 min. Mice were then returned to their home cage for a 2-hour inter-trial interval. During the retrieval (testing) phase, the guillotine door was removed, and mice were given free access to all three arms for 5 min. The number of entries into each arm was recorded. Time spent in each arm was also recorded. The chamber was cleaned with 70% ethanol between subjects and cleaned twice between sex cohorts.

### Statistics

Statistical analyses were performed using GraphPad Prism (version 10.6.1). Statistical tests were selected based on the number of comparisons and experimental design, with specific tests detailed in the respective figure legends. One-way ANOVAs were used to assess the effect of a single treatment variable. Two-way ANOVAs were used to assess the effects of treatment and time. Three-way ANOVAs were used to assess the effects of treatment, time, and sex. Repeated measures ANOVAs were applied when the same subjects were measured across multiple timepoints. Paired t-tests were used for within-animal comparisons (NT vs. *Cdk5-* sgRNA). Normality was assessed using F-tests of variance, and outliers were identified and removed using Grubbs’ or ROUT tests. Post hoc analysis was performed via Tukey’s multiple comparison testing. Statistical significance was set at p < 0.05. All data are presented as mean ± standard error of the mean (SEM).

## Acknowledgments

We would like to thank the Neurobehavioral Testing Core at the University of Pennsylvania for training and equipment support for contextual fear conditioning behavioral assays.

We are grateful to Peter Hamilton at Virginia Commonwealth University and Rachel Neve at University of Massachusetts Medical School for generously sharing CRISPR vectors that were instrumental in the development of these tools.

This work was funded by the National Institutes of Drug Abuse Grant No. R01-DA052465 (EAH) and R01-MH126027 (EAH) and the National Science Foundation Graduate Research Fellowship under Grant No. DGE-1845298 (KLR).

## Author Contributions

KLR and EAH conceptualized and designed the study.

KLR, JJW and AS conducted molecular construct development.

KLR, MA, JJW, and AS performed stereotaxic surgeries.

KLR, AS and KC performed sample collection and preparation.

KLR and MA conducted behavioral assays and molecular analyses.

KLR wrote the original manuscript draft.

KLR, EAH and JJW revised and edited the manuscript.

## Competing Interests

The authors declare no competing interests.

## Data Availability

The CUT&TAG-seq data generated in this study have been deposited in the NCBI Gene Expression Omnibus (GEO) under accession number GSE327281.

## Notes

### Competing Interest Statement

The authors have declared no competing interest.

