## Supplemental Tables and Figures for "Neuron-specific epigenetic repression of *Cdk5* impairs hippocampal-dependent memory in male and female mice"

**Supplemental Material**

**Supplementary Table 1:** qCHIP primers targeting the *Cdk5* promoter.

| Primer | Pair Name | Strand | Start | End | Sequence (5'→3') |
|---|---|---|---|---|---|
| +68F | +68 | − | chr5:24423542 | chr5:24423561 | AGTACCACCTCCTCTGCAAC |
| +68R | +68 | + | chr5:24423362 | chr5:24423381 | GCCGGTATAGCTGACGGTAA |
| −38F | −38 | − | chr5:24423656 | chr5:24423675 | CTTTGTAGTCCGCGTGTCCT |
| −38R | −38 | + | chr5:24423461 | chr5:24423483 | TCTCCAGTTTCTCGTATTTCTGC |
| -228F | -228 | − | chr5:24423835 | chr5:24423855 | AGGCTGTGAGCACAGAAAAGG |
| -228R | -228 | + | chr5:24423660 | chr5:24423680 | CACGCGGACTACAAAGTCCAA |
| -407F | -407 | − | chr5:24423973 | chr5:24423992 | AACACCCAACCAGGTCAGAG |
| -407R | -407 | + | chr5:24423882 | chr5:24423901 | CGCGTTCCAGAATACAGTGA |
| -588F | -588 | − | chr5:24424055 | chr5:24424074 | GTGACCCCAGGACTGATTGG |
| -588R | -588 | + | chr5:24424161 | chr5:24424180 | GTTAGAGTCTGGATTTGAGT |
| -683F | -683 | − | chr5:24424156 | chr5:24424175 | GCCCCGTTAGAGTCTGGATT |
| -683R | -683 | + | chr5:24424250 | chr5:24424269 | ATGTCAAAGTGGAGCCGCTG |

**Supplementary Table 2:** qPCR Primers

| Primer | Sequence (5'→3') |
|---|---|
| *Cdk5 F* | GCTGCCAGACTATAAGCCCTAC |
| *Cdk5 R* | TGGGGGACAGAAGTCAGAGAA |
| *Slc4a2 F* | GGCGCAGATTCTTTGCACAC |
| *Slc4a2 R* | TCCACACCTAAGGTACGAAGTT |
| *Gapdh F* | AGGTCGGTGTGAACGGATTTG |
| *Gapdh R* | TGTAGACCATGTAGTTGAGGTCA |
| *Rpl13a F* | ATGACAAGAAAAAGCGGATG |
| *Rpl13a R* | CTTTTCTGCCTGTTTCCGTA |
| *dCas9-HDAC3 F* | TGAGCATGCCCCAAGTGAAT |
| *dCas9-HDAC3 R* | ACCACCAGCACAGAATAGGC |

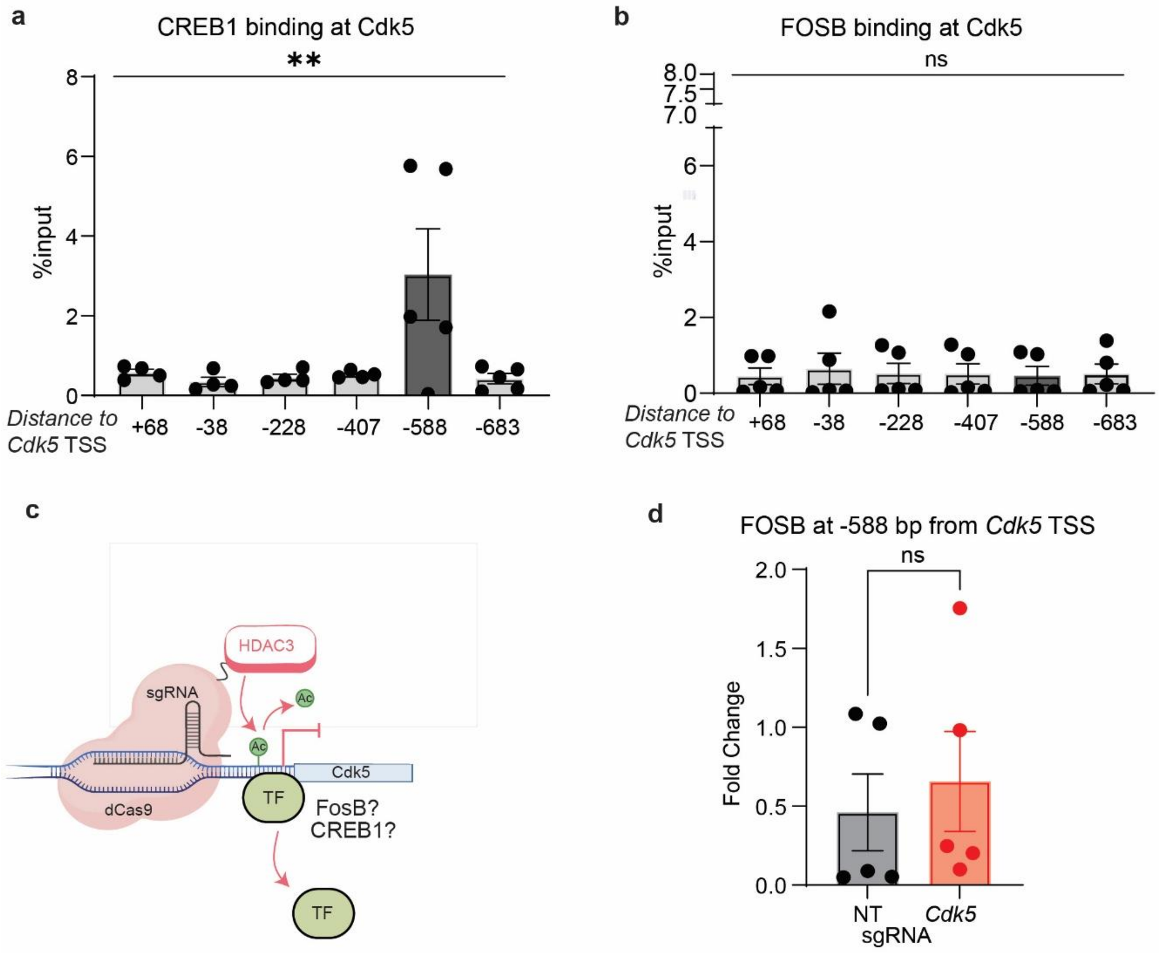


**Fig. S2 CREB1, but not FOSB, is enriched at the *Cdk5* promoter and is displaced by histone deacetylation. (a)** CREB1 qChIP signal across six primer pairs in NT hippocampal tissue shows peak enrichment at ~−588 bp from the *Cdk5* TSS. One-way ANOVA: $F(5, 20) = 3.956$, $p = 0.0115$; post-hoc comparisons at −588 bp vs. +68 bp, $p = 0.0492$; vs. −38 bp, $p = 0.0261$; vs. −228 bp, $p = 0.0353$; vs. −407 bp, $p = 0.0425$; vs. −683 bp, $p = 0.0212$. **(b)** FOSB binding is not enriched at any primer location in NT hippocampal tissue (one-way ANOVA: $F(5, 24) = 0.0597$, $p = 0.9974$, ns). **(c)** Schematic of the experimental hypothesis: transcription factors sharing a predicted binding motif near the *Cdk5* promoter may be regulated by local H3K9/14ac levels. **(d)** FOSB enrichment at −588 bp is unchanged in animals treated with hSyn-dCas9-HDAC3 and *Cdk5*-sgRNA ($n = 5$). All data are presented as mean ± SEM.

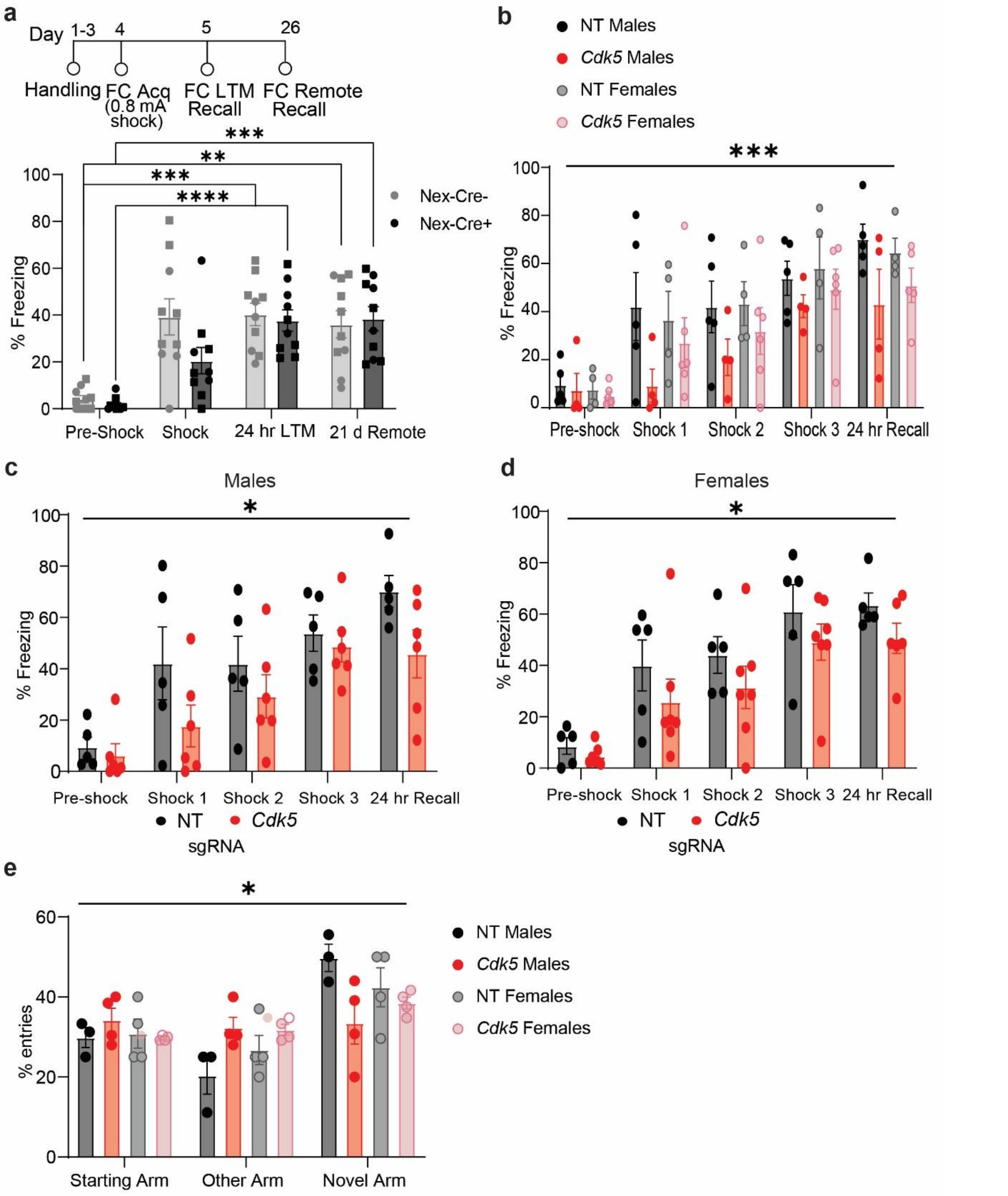


**Fig. S3 Nex-Cre+ mice perform normally in contextual fear conditioning and histone deacetylation at Cdk5 reduces freezing behavior in both males and females. (a)** Contextual fear conditioning protocol performed as described previously[76]. Animals received a single 0.8 mA footshock, and freezing was assessed at 24 hours (long-term memory, LTM) and 21 days (remote memory) post-conditioning. NEX-Cre+ and NEX-Cre− animals exhibit comparable freezing across all timepoints, validating the Cre driver line. Two-way ANOVA revealed a main effect of time ($F(2.441, 43.94) = 32.66$, $p<0.0001$) but no effect of genotype

($F(1, 18) = 1.185$, $p=0.2907$) or genotype × time interaction ($F(2.441, 43.94) = 2.481$, $p=0.0848$). **(b)** Epigenetic repression of Cdk5 in hippocampal excitatory neurons impairs fear memory similarly in both males and females. Three-way ANOVA revealed main effects of time ($F(4, 74) = 18.96$, $p<0.0001$) and sgRNA ($F(1, 74) = 12.17$, $p=0.0008$) but no effect of sex ($F(1, 74) = 0.7115$, $p=0.4017$, ns) or interaction of sex x sgRNA ($F(1, 74) = 1.456$, $p=0.2315$, ns) or sex x time ($F(4, 74) = 0.1728$, $p=0.9517$, $n = 9-10$). **(c)** In males, Two-way ANOVA revealed main effects of time ($F(4, 45) = 11.23$, $p<0.0001$) and sgRNA ($F(1, 45) = 7.081$, $p=0.0108$. $n = 5$), with post-hoc comparisons showing significant differences between NT and *Cdk5*-sgRNA groups at shock 1 ($p=0.0434$) and 24-hour LTM ($p=0.0435$). **(d)** In females, Two-way ANOVA revealed main effects of time ($F(4, 49) = 15.53$, $p<0.0001$) and sgRNA ($F(1, 49) = 5.785$, $p=0.0200$, $n = 5$). **(e)** No sex differences in novel arm preference in the Y-maze. Three-way ANOVA revealed a main effect of arm ($F(1.557, 17.13) = 10.93$, $p=0.0016$) and a significant arm × sgRNA interaction ($F(1.557, 17.13) = 5.112$, $p=0.0243$), but no effect of sex ($F(1, 11) = 0.5011$, $p=0.4937$) or sex × sgRNA ($F(1.557, 17.13) = 0.3836$, $p=0.6570$) or sex × arm interactions ($F(1.557, 17.13) = 0.3836$, $p=0.6359$, $n = 4$).

**Western Blot Raw Images**
**Fig 1f**
Cdk5

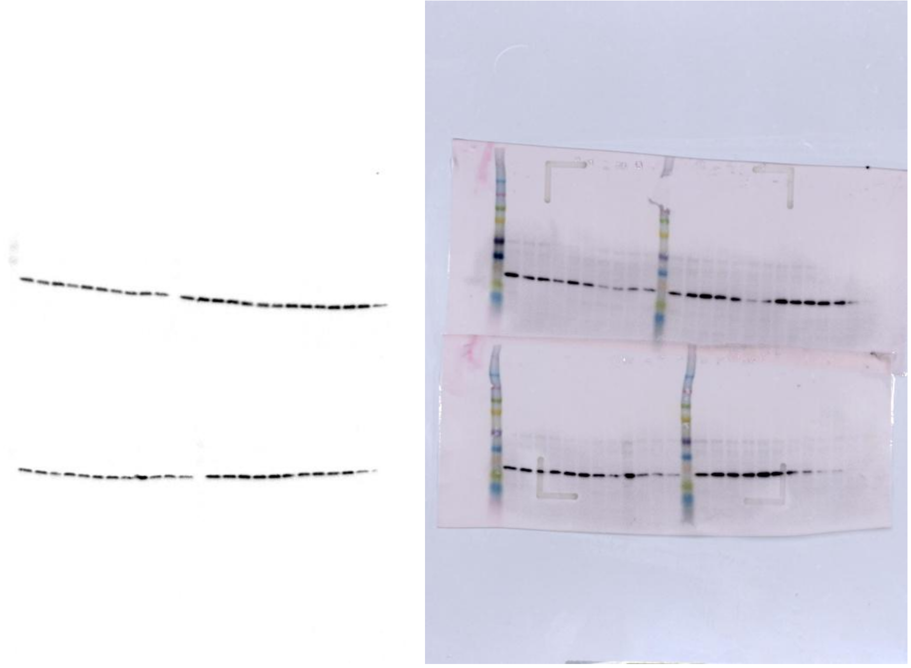

*membrane top-left is 6 days PI
top-right is 14 days PI, bottom-left is 8 days PI and top-right is 9 days PI- data not included in ms

A-tubulin (loading control)

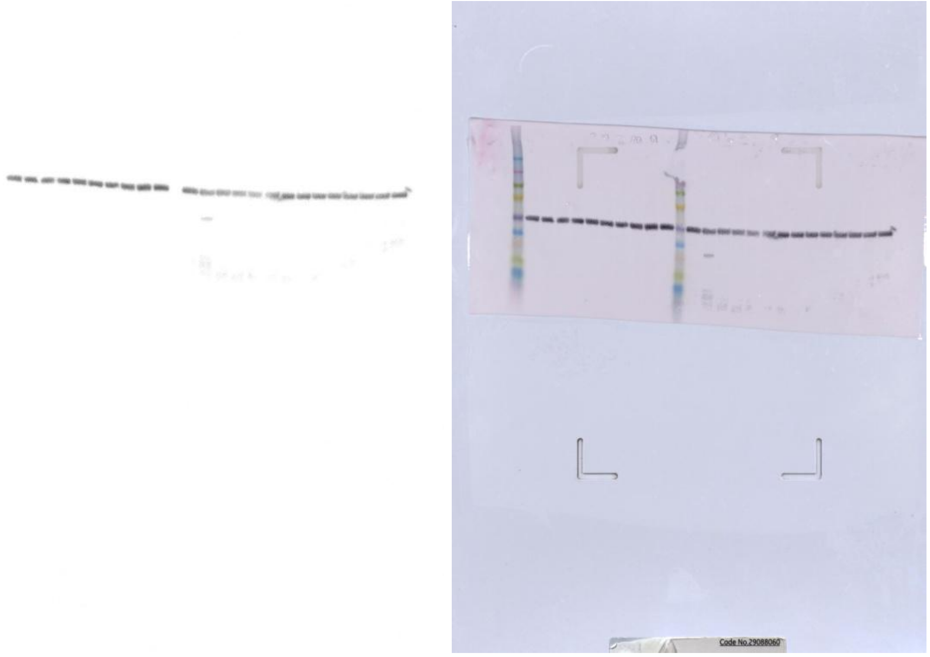

*membrane left is 6 days PI, right is 14 days PI (data not included in ms)

**Fig 1g**

Phospho-Tau(S396)

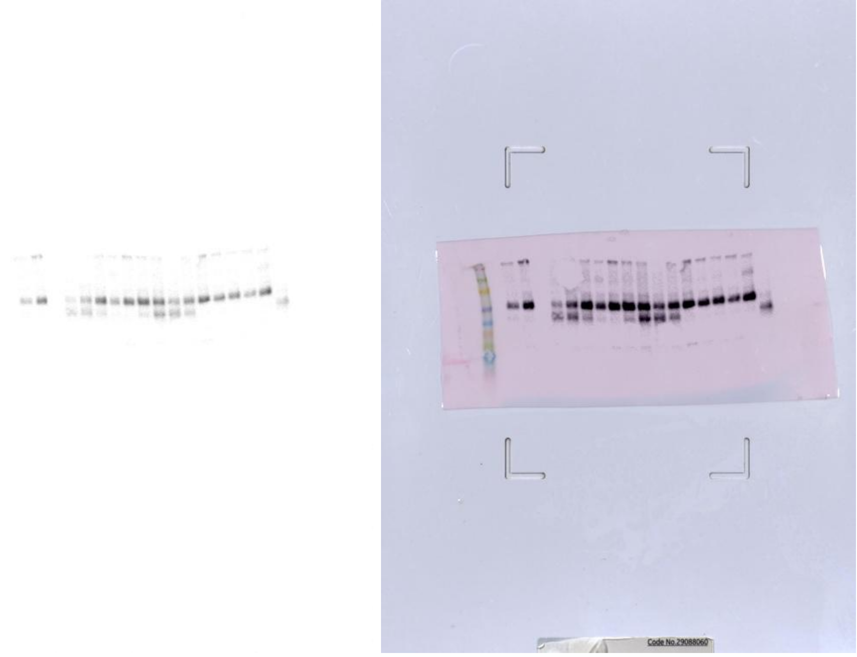

Total Tau

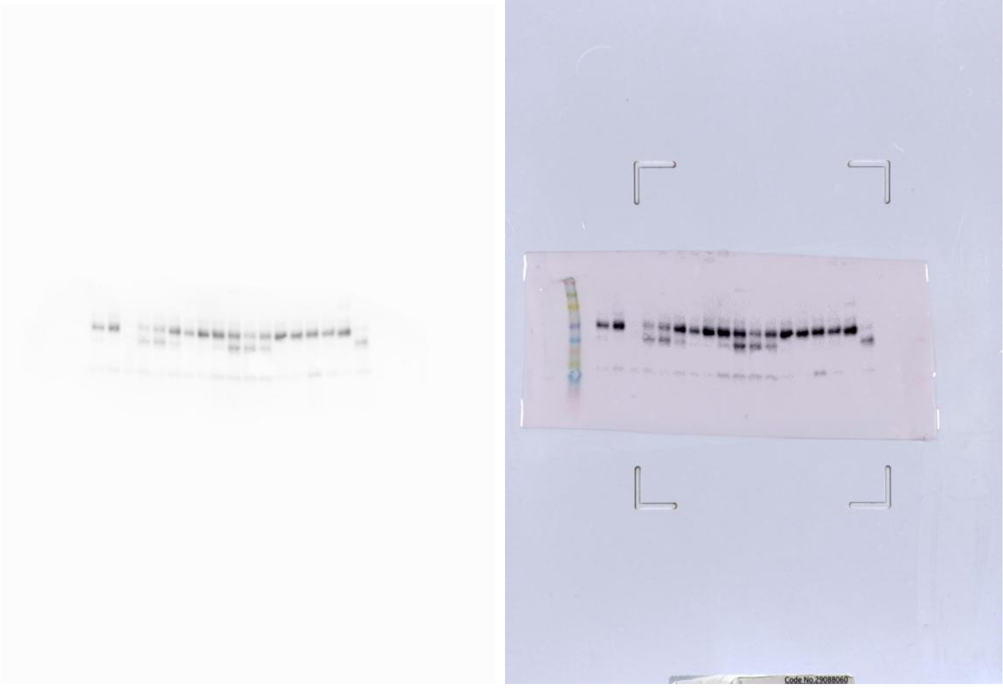

A-tubulin (loading control)

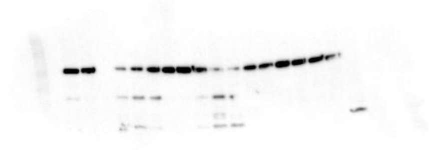

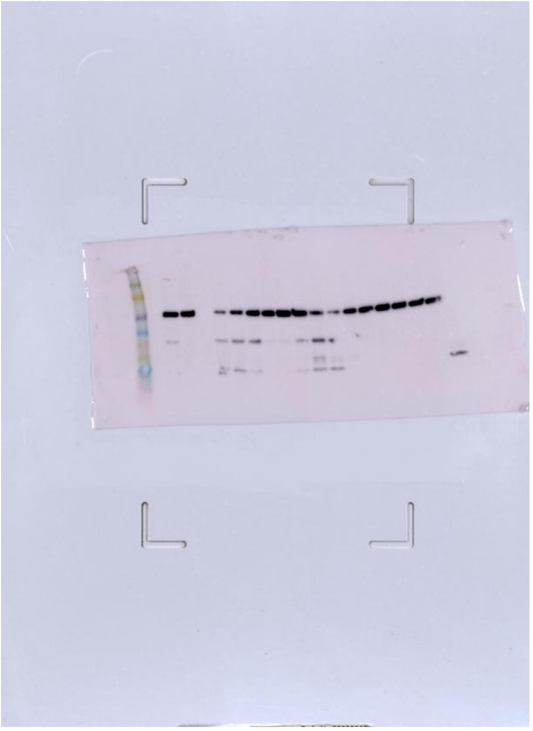
